# The interaction between elapsed time and decision accuracy differs between humans and rats

**DOI:** 10.1101/726703

**Authors:** Carly A Shevinsky, Pamela Reinagel

## Abstract

A stochastic visual motion discrimination task is widely used to study rapid decision-making in humans and animals. Among trials of the same sensory difficulty within a block of fixed decision strategy, humans and monkeys are widely reported to make more errors in the individual trials with longer reaction times. This finding has posed a challenge for the drift-diffusion model of sensory decision-making, which in its basic form predicts that errors and correct responses should have the same reaction time distributions. We previously reported that rats also violate this model prediction, but in the opposite direction: for rats, motion discrimination accuracy was highest in the trials with the longest reaction times. To rule out task differences as the cause of our divergent finding in rats, the present study tested humans and rats using the same task and analyzed their data identically. We confirmed that rats’ accuracy increased with reaction time, whereas humans’ accuracy decreased with reaction time in the same task. These results were further verified using a new temporally-local analysis method, ruling out that the observed trend was an artifact of non-stationarity in the data of either species. The main effect was found whether the signal strength (motion coherence) was varied in randomly interleaved trials or held constant within a block. The magnitude of the effects increased with motion coherence. These results provide new constraints useful for refining and discriminating among the many alternative mathematical theories of decision-making.

## Introduction

The accuracy and timing of sensory discriminations have been used to study mechanisms of decision-making for over a century (*1*). In particular, a visual random dot coherent motion task (*2, 3*) has been extensively used to study decision-making in human and monkeys (*4–22*). In each trial, many small high-contrast dots are plotted at random locations in part of the visual field of a subject. A fraction of the dots move in a coherent direction (“signal”) while others move at random (“noise”). The direction of the coherent motion provides information regarding which of two (or more) available actions will be associated with reward and which associated with non-reward or penalty in the current trial. In the pure reaction-time version of the task, both the time of response and the response selected are freely determined by the subject.

As stimulus strength (signal-to-noise ratio) increases, motion discrimination accuracy increases and reaction time decreases for both monkeys (*23*) and humans (*10*). Instructions that induce the subject to be more conservative result in longer reaction times and higher accuracy. Both these findings are predicted by biased random walk or bounded drift diffusion models (*24*), which postulate that noisy sensory evidence is integrated over time until the accumulated evidence reaches a decision threshold. In this framework, weak stimuli (i.e. with low signal-to-noise ratio) have low drift rates, and therefore accumulate evidence more slowly, reach threshold later, and have a higher likelihood of reaching the wrong threshold first (errors). This parsimoniously explains the psychometric curve (dependence of accuracy on signal strength), chronometric curve (dependence of reaction time on signal strength), and the characteristic long-tailed distribution of reaction times. When subjects are cued to be conservative, this is modeled by setting higher decision thresholds, such that it takes longer for accumulating evidence to reach threshold, but the probability of error is reduced. This provides an elegant explanation of the well-known Speed-Accuracy Trade-Off.

Reaction time and accuracy interact in yet a third way, however. Among trials of equal sensory difficulty tested within a block of fixed decision criterion, studies of humans and other primates widely report that the trials with longer reaction times are more likely to be errors. This is not predicted by the simple bounded drift-diffusion model outlined above, which predicts no correlation between reaction time and accuracy. Several variants of the drift-diffusion model can account for the result, however, for example by postulating variable drift rates, collapsing bounds, or urgency signals (reviewed in (*1, 25, 26*)).

We previously trained rats to perform random dot motion discrimination and characterized the speed and accuracy of their decisions (*27*). Rats’ accuracy was dependent on the duration of stimulus presentation, and the increase in accuracy with reaction time was contingent on stimulus presence. Moreover, reaction time declined with stimulus strength, and the shape of the reaction time distribution resembled the characteristic distribution produced by a bounded diffusion process. These results are consistent with the hypothesis that rats are accumulating evidence in the task. The data from rats differed from the primate literature in one key respect: in that study, rats’ later decisions were more likely to be accurate. This result has important implications, because some of the model modifications proposed to explain the primate data would be incompatible with the rat data.

Given the apparent inconsistency of the result with the past literature, however, replication and further characterization of the result remained important. It was unclear whether the difference reflected a true species difference, or whether it arose from differences in the trial structure, task implementation or data analysis. Furthermore, the potential for artifacts due to non-stationarity has been under-appreciated in the literature. Undetected behavioral non-stationarity is rather likely, and might have caused spurious correlations between accuracy and reaction time in the data of either or both species. The experiments and analyses presented here address these concerns, ultimately confirming that the dependence of accuracy on reaction time is different between species, and not an artefact of non-stationarity in either case.

## Results

### Experimental approach

Our previously reported experiments with rats (*27*) used a similar visual random-dot coherent motion stimulus as the published primate studies to which we compared, but the trial structure was quite different. Notably, the rat experiments had no delay between trial engagement and stimulus onset, no delay between a correct response and reward delivery, and no enforced inter-trial interval. Experiments in the primate literature often enforced random delays to stimulus onset, delays to reward delivery, and/or fixed inter-trial intervals. Such task differences affect the anticipation of sensory and reward events and the opportunity cost difference between early and late responses. We also implemented the task in our own custom software and hardware, which presumably differed in numerous details from those used in other studies. Thus it remained possible that task differences rather than species differences caused the difference in results.

To rule this out, in the present study both humans and rats were tested in the same task, with identical trial structure using the same hardware and software. The visual stimulus was a field of randomly located, moving white squares (“dots”) rendered on a black background. A fraction of the dots (“signal”) moved coherently in the same direction, either toward the left or right. The remaining dots (“noise”) were relocated randomly each video frame. The coherence (fraction of dots participating in signal) was either varied in randomly interleaved trials, or held constant within a testing session. The task structure was two-alternative forced-choice: the subject was rewarded for selecting the response on the side toward which the coherent motion flowed. Correct (rewarded) responses were immediately indicated by an audible beep. Errors were penalized by a fixed time-out delay (1-2 s) before a new trial could be initiated. There was no deadline to respond, but most responses were between 500-2500 ms and reaction times exceeding five seconds were rare in both species. Additional task details, including the remaining task differences between species, are described in Methods.

### Visual Performance

For the purpose of studying perceptual decision-making, the visual signal must be perceptible but challenging. Humans and rats have different optics and visual systems. Therefore we used different display parameters (viewing distance, spatial extent of the visual motion field, and the number, contrast, size, and speed of the moving dots) and different motion coherence ranges for humans and rats, although we did not attempt to optimize parameters in either case. We analyzed visual performance in blocks of trials in “epochs” during which stimulus and reinforcement parameters were fixed and subject performance was not trending (discussed in more detail below).

Motion coherence threshold is likely to be a function of other stimulus parameters, such as dot size, dot density, contrast, and motion speed. Since these parameters were not varied parametrically or optimized for either species, we cannot interpret the observed motion threshold as an absolute measure of motion perception ability in either species. The dot size, dot density, contrast and motion speed we chose for humans resulted in lower motion thresholds and higher motion sensitivity in human subjects than the parameters we chose for rats produced in rats (Fig. 1 A-F vs. G-L). Nevertheless, with the stimulus parameters we used, the overall reaction times of rats and humans were similar, with a minimum response latency of ~0.5 s, a median of ~1 s, and 95^th^ percentile of ~3s in both species (not shown). Motion discrimination performance ranged from 50% (chance) to 100% as the motion coherence increased for most subjects.

**Figure 1.**
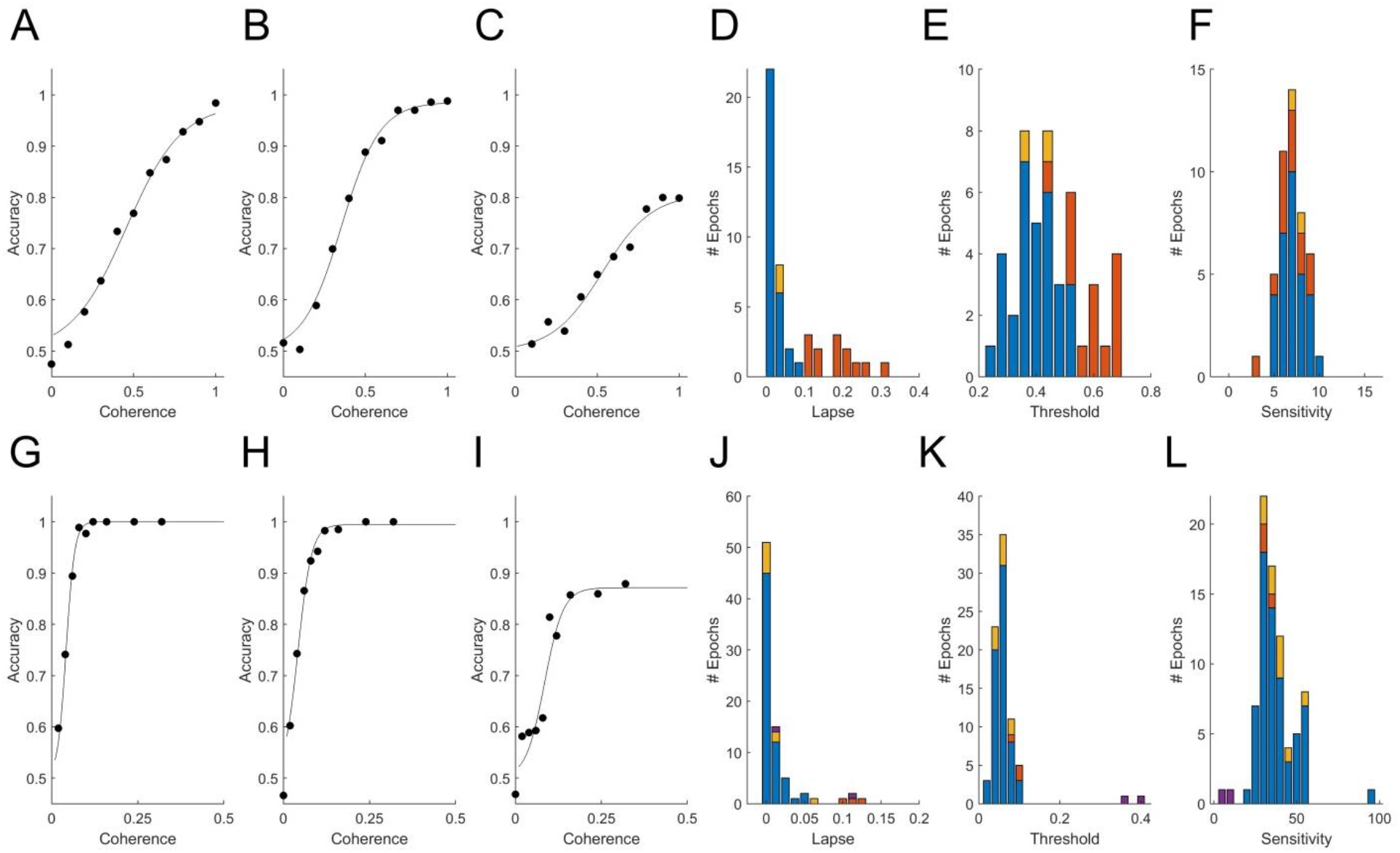
Psychometric functions of rat and human populations. **A-C** and **G-I**: Accuracy (fraction correct) as a function of stimulus strength (coherence) for three rats (A-C) and three humans (G-I) from our data set. Points indicate fraction correct among trials at the indicated coherence. Curves are psychometric fits, from which we obtain lapse (extrapolated error rate at 100% coherence), threshold (coherence at 75% correct), and sensitivity (slope of psychometric curve) for each epoch (see Methods). Panels **D, J**: Distribution of lapse among qualifying epochs of rats and humans respectively (see Methods for inclusion criteria). Panels **E, K**: Distribution of motion discrimination threshold. Panels **F, L**: Distribution of motion sensitivity; note different scales. In all histograms in this figure, blue indicates epochs with <10% lapse and no bias; red indicates epochs with >10% lapse (e.g., panel C,I), some of which were also biased; yellow indicates epochs with response bias but not lapse; and purple indicates an atypical visually impaired human subject. The outlier near 100 in panel L is the example shown in panel G.

Results of psychometric fits from all non-trending data epochs with a sufficient coherence range are summarized in Fig. 1 D-F (N=46 epochs from 11 rats) and Fig. 1 J-L (N=79 epochs from 52 humans). Most subjects (7/11 rats and 46/52 humans) had at least one epoch in which accuracy was ≥90% on the strongest stimuli; these epochs are indicated by blue in all Fig. 1 histograms. Epochs with high lapse (>10% lapse, i.e., <90% correct at the highest coherence) were more frequent among rats than humans (red in Fig. 1 histograms). In some cases, performance plateaued as a function of contrast below 90% correct (Fig. 1 C,I), indicating that the rat sometimes responded incorrectly even when stimulus was perceptually obvious. Some but not all of these could be explained by response bias (preference to respond left or right). In other cases of high lapse, however, rats’ psychometric curves were still not leveling off at 100% coherence (not shown). In these cases interpretation of lapse is unclear, as other perceptual factors such as acuity or contrast sensitivity may have been limiting their performance. In all epochs with lapse, measurement of threshold was problematic. If lapse reflected a vertical scaling of the psychometric curve without horizontal shift, the measure of perceptual threshold we used (coherence at which the subject reaches 70% correct) would be artificially high (cf., red in Fig. 1 E,K). Some epochs that did not exhibit lapse still had substantial response bias (shown in yellow in Fig. 1). This occurred because response bias was exhibited only in low-coherence trials – as if the subject relied on a prior only when sensory evidence was weak. Because of these caveats, epochs with high lapse or bias were either excluded in subsequent analyses, or analyzed separately.

In summary, analysis of visual performance confirmed that we have data from both species in which the subjects were under stimulus control and for which performance was perceptually limited.

### Relationship between RT and accuracy confirmed for rats and humans

One way to visualize the species difference is to compare the reaction time distributions of errors to that of correct trials. For rats, we found that the reaction time distributions of correct trials had heavier tails (Fig. 2A), causing a rightward shift of the cumulative probability distribution of correct trials relative to errors (Fig. 2B). Therefore accuracy increased with reaction time (Fig. 2C). For rats in general, the average reaction time of correct trials was greater than that of error trials of the same coherence within the same experimental test block (Fig. 2D). Therefore the accuracy of slow responses was lower than that of fast responses (Fig. 2E). For humans, we found the opposite: reaction times of errors were longer, and therefore accuracy declined with reaction time (Fig 2F-J). Thus we can replicate the result that has been widely reported in past studies of humans and monkeys, using the same trial structure, task implementation, and data analysis procedures that yields the opposite result for rats.

**Figure 2.**
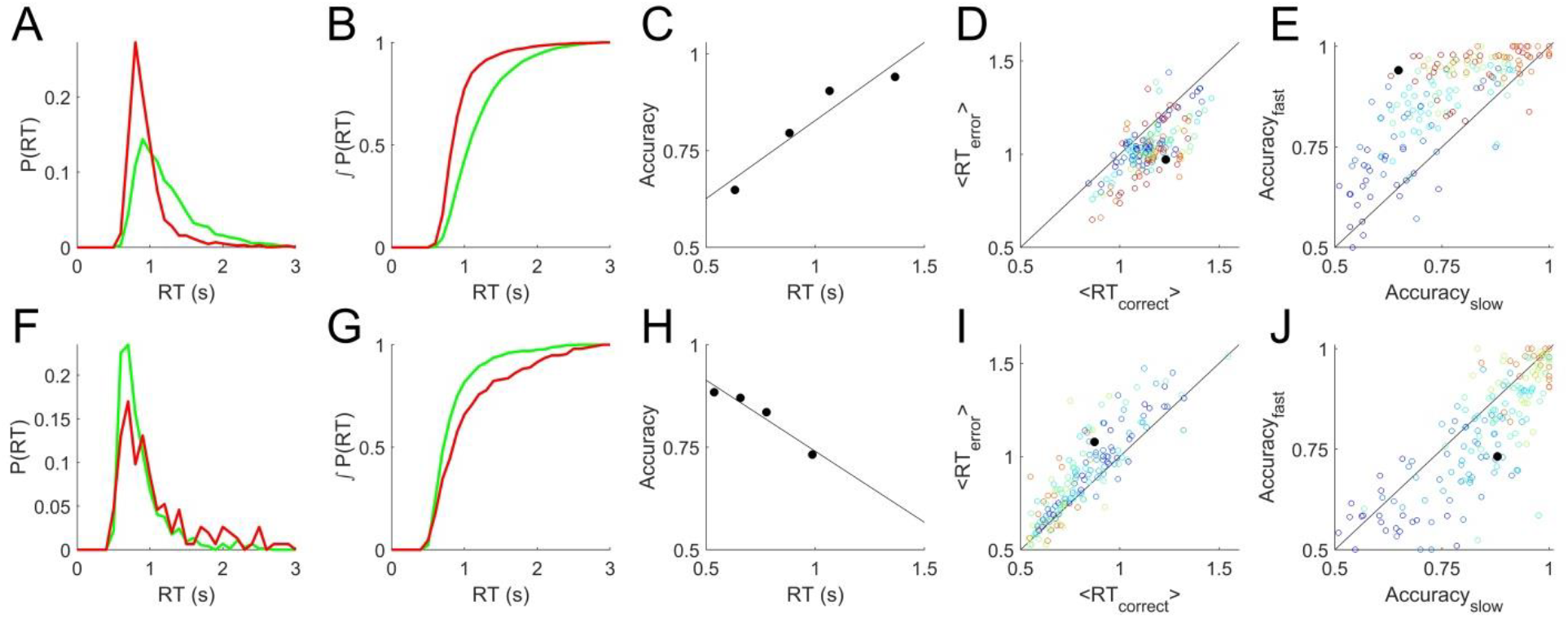
Relationship between reaction time and accuracy. **A.** Example reaction time probability distributions from a rat single-coherence experiment, N=10,858 trials, 82% correct. Green=correct trials, red=error trials. **B.** Cumulative distributions, the integrals of curves in A. **C.** Accuracy vs. reaction time quartiles (symbols) and linear regression to these points (line), for the data shown in A and B. **D.** Average reaction time of correct trials vs. of error trials in rat data. Each symbol represents analysis of all trials of like coherence from one non-trending, unbiased epoch. Colors indicate coherence from 0.1 (blue) to 0.85 (red). Black filled circle represents the data shown in A-C. **E.** Accuracy of slow trials vs. fast trials from rat epochs described in D. Slow and fast defined as the bottom and top quartiles of reaction time respectively for that coherence in that epoch. Symbols as in D. **F.** Example reaction time probability distributions from a human subject from a single-coherence epoch with N=909 trials, 83% correct, colors as in A. **G.** Cumulative distributions, the integrals of curves in F. **H.** Accuracy vs. reaction time for data in F-G. **I.** Average reaction times of correct vs. error trials from human epochs selected as described in D, also excluding the visually impaired human subject. Analyzed as in D; symbols as in D except that coherence scale is 0.02 (blue) to 0.16 (red). **J.** Accuracy of slow vs. fast responses for human experiments, analyzed as described in E, symbols as in I. Additional details in Methods.

To summarize the sign and strength of this trend, we determined the slope of the regression line of accuracy vs. reaction time for all candidate epochs (see Methods for details). Epochs with fixed coherence were analyzed as shown in Fig. 2 C,H. In the epochs with randomly interleaved coherences, there were often not enough trials of a single coherence to reliably estimate the accuracy in multiple reaction-time bins.

Therefore in all psychometric epochs we pooled trials from the coherences for which the subject was between 70-90% correct (c.f. Fig. 1 A-C,G-I) to estimate accuracy vs. reaction time (e.g., Fig. 3A,D). In epochs with sufficient numbers of trials to estimate trends within each coherence, these were consistent with the coherence-pooled slope for the same epoch (e.g., Fig. 3B,E).

**Figure 3.**
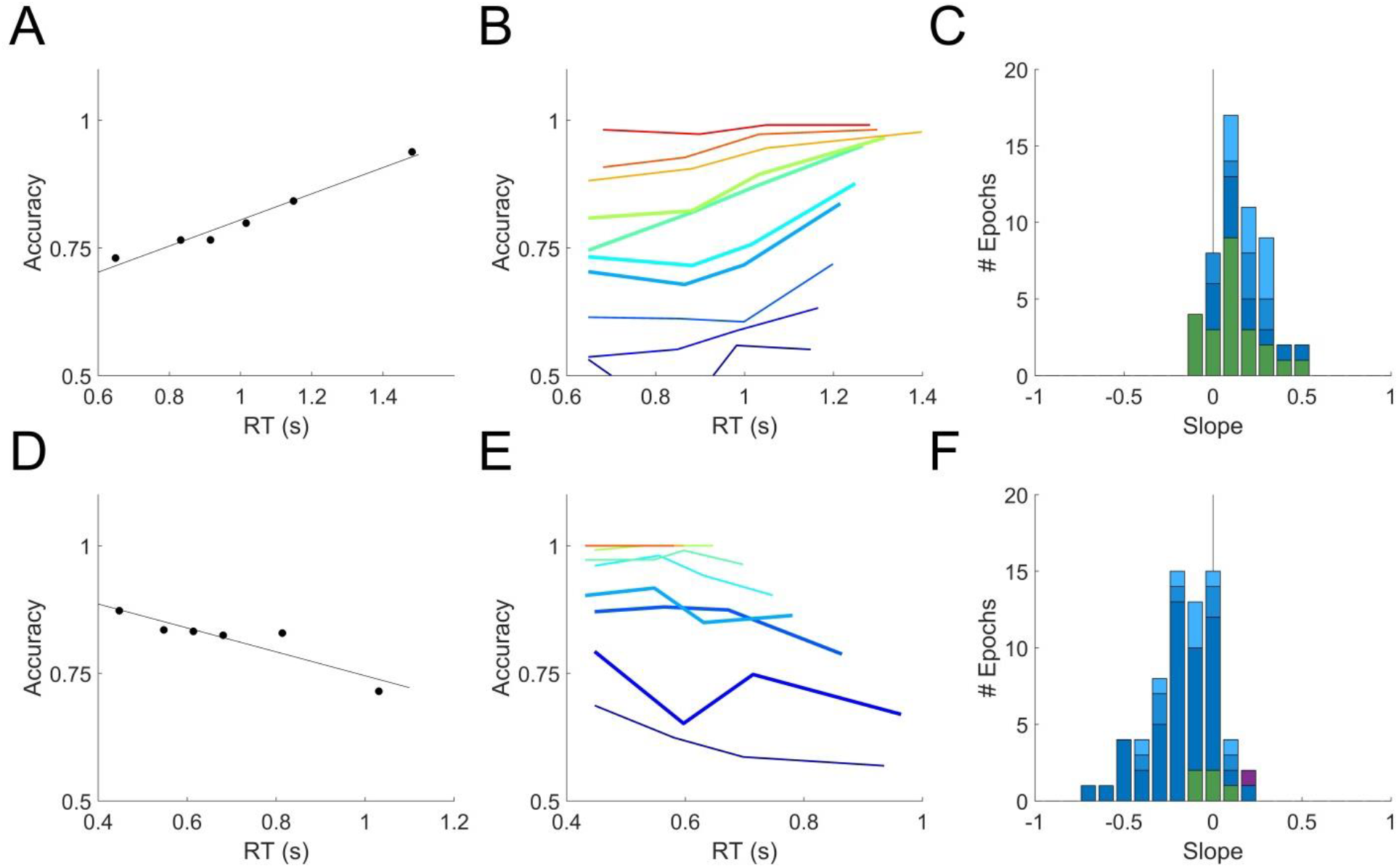
Relation of accuracy to reaction time is opposite in rats and humans. **A.** Dependence of accuracy on reaction time for an example rat subject evaluated within a single epoch for all trials within the sensitive range (70-90% correct) of psychometric curve. Symbols show mean accuracy and mean reaction time, in reaction time bins with equal numbers of trials; the slope of the linear regression (line) was +0.26 for this epoch. **B.** Accuracy vs. reaction time for the same data as in A, computed separately by coherence. Color indicates coherence, 10-100% coherence (blue→red). Thick curves indicate the coherences pooled in panel A. This epoch contained 8149 qualifying trials. **C.** Distribution of slopes computed as in A, for all qualifying rat epochs (see Methods). Colors indicate epochs with <1% lapse (dark blue, N=12), 1-2% lapse (medium blue, N=8), 2-10% lapse (pale blue, N=10), or fixed coherence (green, N=23). **D.** Dependence of accuracy on reaction time for an example human subject within a single epoch, analyzed as described in A. Slope was −0.23 for this epoch. **E.** Data from the example human subject in D, analyzed as in B. Coherences 2-32% (blue→red). This epoch contained 5652 qualifying trials. **F.** Distribution of slopes in all qualifying human epochs: <1% lapse (N=46), 1-2% lapse (N=7), 2-10% lapse (N=8) and fixed coherence (N=5) colors as in C; visually impaired subject (N=1) in purple. Details of analysis are in Methods.

The distribution of slopes for rats (Fig. 3C) and humans (Fig. 3F) are shown for all non-trending, unbiased, low-lapse (if psychometric) epochs. For rats the average slope was 0.16 ± 0.14 overall (mean±SD; N=53 epochs from 11 rats), and 0.18 ± 0.14 in psychometric epochs with <1% lapse (N=12). For humans the average slope was – 0.16 ± 0.18 overall (N=66 epochs from 48 humans, excluding the visually impaired subject), and – 0.19 ± 0.19 in psychometric epochs with <1% lapse (N=46). These data confirm our earlier report that for rats, accuracy increases with reaction time in this task. Moreover, most humans showed the opposite trend: accuracy declined with reaction time, consistent with previous reports of humans and non-human primates on similar tasks. The results were opposite for rats and humans, but of equal magnitude and consistency within either species.

### The problem of non-stationarity

Reaction time distributions are extremely broad; comparing the shapes of the correct and error reaction time distributions, particularly differences in the tails of the distributions, requires a substantial number of error trials for each value of motion coherence evaluated. Therefore analyses like those of Figure 2 and 3 require long experiments, particularly if coherence values are interleaved or for coherences with low error rates. It is crucial to validity of the analysis that the underlying distributions are *stationary*, i.e., not changing over the time these samples are taken to estimate them.

To illustrate why this is important, consider some extreme examples of non-stationarity. Most subjects get faster and more accurate with learning and practice; if this change were occurring over the range of trials included in an analysis, reaction time would be *negatively* correlated with accuracy for uninteresting reasons. Therefore we and others routinely exclude learning period data, analyzing only the trials after performance is deemed stable. Or suppose subjects occasionally and implicitly shift strategies, either increasing or decreasing their threshold at unknown times for an unknown duration. It is known that a cued increase in the decision threshold for a block of trials will increase both reaction time and accuracy in that block, so we would expect such un-cued threshold changes to induce a *positive* correlation between reaction time and accuracy in the aggregated data, even if these variables were uncorrelated or negatively correlated among the trials performed with any given decision threshold. Therefore we and others routinely inspect data for overt signs of non-stationarity, such as long-term trends or abrupt shifts in reaction time distribution or accuracy. Finally suppose a subject’s motivation or attention wandered over the course of a two hour session. Depending on failure mode (which may be species-specific) this could induce positive or negative correlated fluctuations in both accuracy and reaction time.

The ability to detect non-stationarity is statistically limited, and to our knowledge a fully satisfactory method does not exist. Linear trends that are present may not be statistically distinguishable from chance due to limited power. A subject’s state could fluctuate on shorter timescales with no net trend. Indeed we find in both species that if we compute the average reaction time in non-overlapping blocks of 16 trials within an apparently stationary epoch, the variance over blocks is nearly always significantly higher than chance, as measured in time-shuffled controls (not shown). Fluctuations in state are just as problematic as long-term trends for our analysis, but fluctuations on short timescales are even more difficult to detect statistically, and may be impossible to avoid experimentally. Therefore it is important to consider the possibility that either the positive correlation between reaction time and accuracy in rats, or the negative correlation in humans, or both, could arise from undetected non-stationarity.

To address this concern, we developed a complementary method to assess the difference in reaction time between correct and error trials using only temporally local comparisons. The premise of this approach is that the shorter the time window, the more likely the subject’s state is well approximated as stationary. We computed the difference in reaction time between each error trial and the closest correct trial of the same coherence that was at least 3 trials before it, and report the mean of these local time differences. We performed this analysis two ways: either each epoch was analyzed separately (Fig. 4 A-C,G-I), or each subject’s lifetime trial data were analyzed *en masse*, ignoring blatant non-stationarities including initial learning, session warm-up, and changes in task parameters (Fig. 4 D-F,J-L). The similarity of the results of these two analyses shows the robustness of this method to non-stationarity.

**Figure 4.**
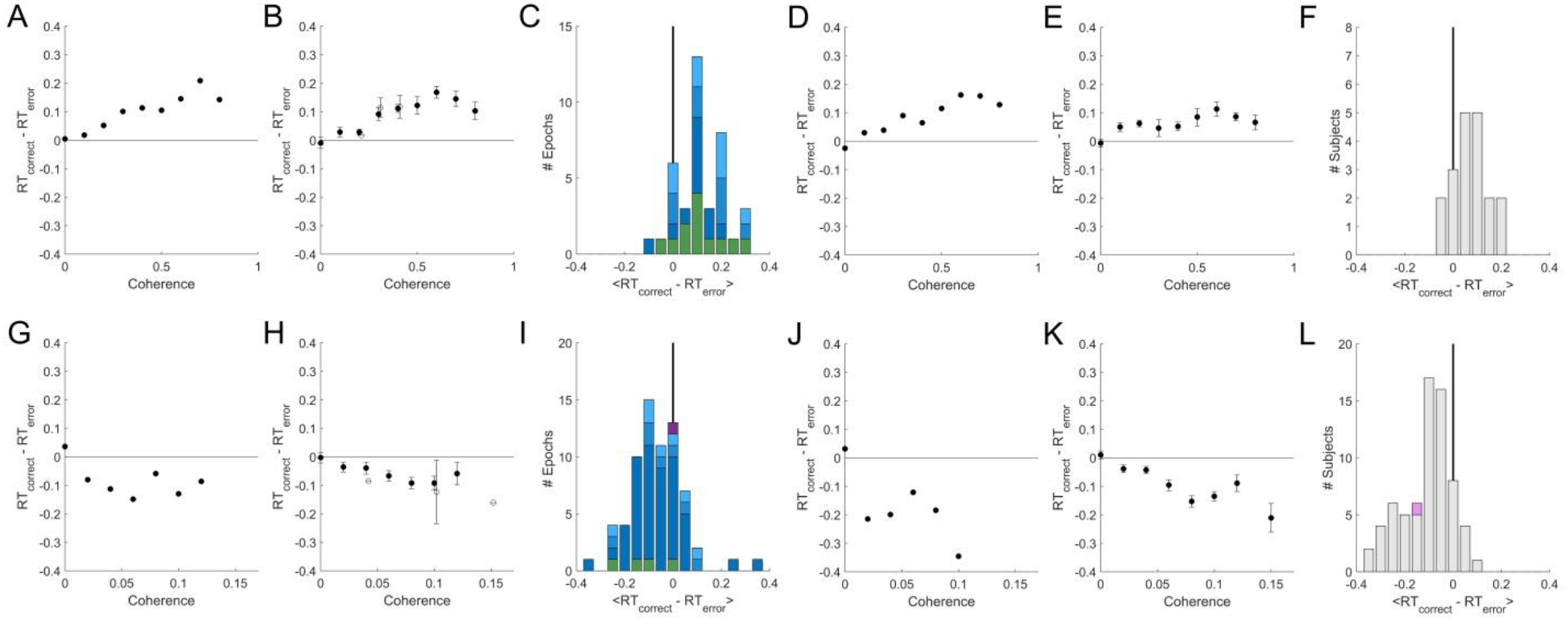
A temporally-local measure of the interaction between accuracy and reaction time. **A.** The mean difference between reaction times of nearby correct and error trials as a function of coherence, for an example epoch from a rat. **B.** Population average of the mean difference in correct vs. error reaction times within coherence, averaged across psychometric epochs (solid) or fixed-coherence epochs (open). Error bars show SEM over the number of epochs for which the value of the respective coherence was estimated. **C.** Distribution over rat epochs of the mean reaction time difference between correct and error trials (computed within coherence as in B, then averaged within epoch over all coherences >=0.4). Colors as defined in Fig. 3C. **D.** Like panel A but computed from the lifetime trial data of one example rat subject, not excluding any trials. **E.** Like panel B, but averaged over lifetime-analyses of all rat subjects. **F.** Like panel C, but distribution is over rat subjects, each estimated from a lifetime analysis. **G-L:** like A-F but for human subjects. **G.** Example human epoch. **H.** Population average over psychometric (solid) and fixed-coherence (open) epochs of non-impaired human subjects. **I.** Distribution of mean reaction time differences all human epochs (averaged within epoch for coherences >=0.04), colors as defined in Fig. 3F. **J.** Example human subject lifetime analysis. **K.** Population average over non-impaired human subjects from lifetime analysis. **L.** Distribution over human subjects based on lifetime analysis; visually impaired subject in purple. See Methods for additional details.

For rat subjects, error trials consistently had shorter reaction times than their nearest correct trial of the same coherence (Fig. 4 A-F), whereas for human subjects error trials consistently had longer reaction times (Fig. 4 G-L). This result was the same whether we compared to the correct trials preceding or following the reference error trial (not shown). In both species, the difference between errors and correct trials was greater for higher coherence stimuli. This trend was found in psychometric epochs regardless of lapse (Fig. 4C,I, shades of blue) and in most epochs that used a fixed coherence (Fig. 4 C,I green). In conclusion, although non-stationarity cannot be fully eliminated from the data, such non-stationarity does not trivially explain the dependence of accuracy on reaction time in either humans or rats, nor the difference between the species.

## Discussion

This study confirms our earlier report that for rats performing random dot motion discrimination, errors have faster reaction times than correct trials (Fig 2A-B), and therefore rats have higher accuracy at longer reaction times (Fig. 2C; 3A-C), when comparing trials of like coherence within testing blocks of apparently stable decision threshold. We also demonstrate that in our task, human errors have longer reaction times than correct trials (Fig 2D-E) and thus humans have lower accuracy at longer reaction times (Fig. 2F; 3D-F). By using the same task for both species, we can now rule out the possibility that task differences (which were substantial) accounted for the difference between our previously reported finding in rats and the typical finding in humans and other primates.

An increase in accuracy with reaction time has also been observed in image discrimination by rats (*28*) and in orientation discrimination by rats (our unpub. obs.) and mice (*29*). This trend may have escaped notice in other studies because rodent tasks often use go-no-go designs or cued response times, whereas this analysis requires a reaction time task, in which response time is dictated by the animal and distinct from choice.

One visually impaired human subject was tested with customized stimuli that yielded a motion coherence threshold and sensitivity comparable to what we found in rats. While it would require much deeper investigation, it might not be a coincidence that the relationship between reaction time and accuracy in this subject was similar to that of rats. The subject self-reported to have normal or corrected-to-normal vision, and did not present evident visual disability during consenting and task instruction procedures. We lack data to determine, however, whether this subject’s impairment was specific to motion discrimination or secondary to an impairment in contrast sensitivity, acuity, or motion speed sensitivity. We speculate that it does not matter which stimulus dimension is limiting for performance, but this assumption could be tested. To explore this further one would have to parametrically vary all these stimulus parameters in each subject, and obtain psychometric curves of each parameter dimension using peak or above-ceiling values for all other parameters.

One interpretation might be that subjects for whom the visual task is extremely difficult employ different strategies, regardless of species. We note however that all subjects were assessed using stimuli that were comparably difficult for that subject, as judged by their accuracy (~80%) and comparable mean reaction times. In each subject we know performance was perceptually limited because accuracy increased with coherence, and the effects we report were found at coherences above chance and below ceiling. We also note that the difference between error and correct reaction times increased with coherence. This suggests that making the task easier for the rat does not tend towards producing a more human-like behavioral pattern.

We emphasize that our claim is distinct from two other phenomena that are often confused and could be confounded. First, subjects make faster decisions about stronger sensory stimuli. In bounded diffusion models of decision making, this fact is attributed to different *rates* of evidence accumulation. Correlation between accuracy and reaction time in this sense appears to be the same for rodents and primates, and is not the phenomenon of interest. If one analyzed a mixture of low-coherence and high-coherence trials, however, the early and late decisions would differ in the proportion of trials with low vs. high coherence stimuli, muddling the interpretation of any relationship between average accuracy and reaction time. To avoid this confound, when analyzing experiments that had interleaved coherences we only compared accuracy vs. reaction time within a narrow band of coherence. Nevertheless in psychometric experiments it was usually necessary to pool over a few similar coherences to get enough trials to resolve the accuracy as a function of reaction time (e.g., in Fig. 3 A,D). In the epochs that had enough trials, however, we confirmed that the trend held within each discrete coherence (e.g., Fig. 2B,E). Moreover, this caveat does not apply to the fixed-coherence epochs, and this subset of epochs had qualitatively similar results (Table 1).

**Table 1.** Analysis of slopes of accuracy vs. reaction time in different subsets of epochs. Indented pairs of lines indicate two mutually exclusive and exhaustive subsets of the parent category. All groupings that were analyzed are shown. Shaded cells indicate exceptions to the main reported effect. The number of epochs and the number of unique individuals included in the subset is indicated by “N eps” and “N subj” respectively. “Trials per epoch” indicates the average number of trials contained in the selected reaction time bins for the regression of binned accuracy vs. binned reaction time. The absolute number of epochs with a negative slope (out of N eps) is given in “# eps slope <0”. The mean and standard deviation of slopes within the subset are shown in “Slope (mean±SD)” The subsets in the bottom 6 rows (*) correspond to the histograms in Figure 3.

|  | Rats |  |  |  |  | Humans |  |  |  |  |
| --- | --- | --- | --- | --- | --- | --- | --- | --- | --- | --- |
| Subset | N<br>eps | N<br>subj | Trials<br>per<br>epoch | # eps<br>slope<br><0 | Slope<br>(mean±S<br>D) | N<br>eps | N<br>subj | Trials<br>per<br>epoch | # eps<br>slope<br><0 | Slope<br>(mean±S<br>D) |
| All candidate epochs | 111 | 18 | 2411 | 14 | 0.20±0.17 | 102 | 62 | 1457 | 81 | -0.15±0.20 |
| All Non-trending epochs | 87 | 17 | 2261 | 9 | 0.20±0.17 | 81 | 56 | 1525 | 66 | -0.15±0.18 |
| All Trending epochs | 24 | 13 | 2953 | 5 | 0.16±0.17 | 21 | 19 | 1193 | 15 | -0.15±0.24 |
| Among Non-trending |  |  |  |  |  |  |  |  |  |  |
| Psychometric | 43 | 11 | 2257 | 1 | 0.23±0.14 | 74 | 52 | 1552 | 61 | -0.15±0.19 |
| Fixed Coherence | 44 | 13 | 2265 | 8 | 0.18±0.19 | 7 | 6 | 1240 | 5 | -0.08±0.13 |
| Epochs with ≥1K trials | 66 | 15 | 2742 | 8 | 0.20±0.17 | 57 | 44 | 1845 | 50 | -0.15±0.17 |
| Epochs with <1K trials | 21 | 11 | 749 | 1 | 0.22±0.18 | 24 | 22 | 767 | 16 | -0.13±0.21 |
| Psycho >10% lapse | 11 | 4 | 2596 | 0 | 0.36±0.16 | 3 | 2 | 651 | 1 | 0.00±0.34 |
| Psycho ≤10% lapse | 32 | 7 | 2141 | 1 | 0.19±0.11 | 71 | 51 | 1590 | 60 | -0.16±0.18 |
| Humans Phase 1 |  |  |  |  |  | 11 | 7 | 1461 | 8 | -0.06±0.14 |
| Humans Phase 2 |  |  |  |  |  | 70 | 49 | 1535 | 58 | -0.16±0.19 |
| Rats 2hr sessions | 50 | 16 | 1729 | 7 | 0.21±0.19 |  |  |  |  |  |
| Rats 24hr sessions | 37 | 9 | 2981 | 2 | 0.20±0.15 |  |  |  |  |  |
| Biased | 26 | 13 | 2973 | 3 | 0.25±0.20 | 14 | 12 | 1040 | 11 | -0.10±0.18 |
| Unbiased | 61 | 14 | 1958 | 6 | 0.19±0.16 | 67 | 49 | 1627 | 55 | -0.16±0.18 |
| *psycho <10% lapse,<br>visually impaired |  |  |  |  |  | 1 | 1 | 1353 | 0 | 0.24±0.00 |
| psycho <10% lapse | 30 | 7 | 2170 | 1 | 0.19±0.12 | 61 | 45 | 1660 | 52 | -0.18±0.18 |
| * 0-1% Lapse | 12 | 7 | 1762 | 0 | 0.18±0.14 | 46 | 36 | 1867 | 40 | -0.19±0.19 |
| * 1-2% Lapse | 8 | 4 | 2737 | 1 | 0.17±0.10 | 7 | 7 | 1188 | 5 | -0.15±0.19 |
| * 2-10% Lapse | 10 | 4 | 2207 | 0 | 0.22±0.09 | 8 | 7 | 878 | 7 | -0.14±0.15 |
| *fixed coherence | 23 | 9 | 1552 | 5 | 0.12±0.16 | 5 | 4 | 1280 | 3 | -0.02±0.08 |

The second distinct phenomenon is that when subjects try harder to avoid errors, they tend to take more time to respond (higher average reaction time), and are also more accurate – the well-known Speed-Accuracy Trade-Off. In the bounded diffusion framework, correlations between accuracy and reaction time in this sense are attributed to differing *decision bounds* in testing blocks with differing priority instructions or reward contingencies. Correlation between accuracy and reaction time in this sense appears to be the same for rodents and primates, and is not the phenomenon of interest. But if one unknowingly combined data from two time-ranges that differed in decision criterion, this would introduce a positive correlation between reaction time and accuracy, even if the opposite relationship held among the trials within any narrower time window. To avoid this potential confound, we analyzed data within epochs that used constant testing conditions (stimulus and reinforcement parameters), and that lacked detectable trends in performance (accuracy or reaction time).

Nevertheless non-stationarity may be impossible to detect or avoid. Even if statistical tests fail to detect significant non-stationarity in a time series, such tests have very limited statistical power, only test for specific kinds of non-stationarity, and make assumptions about the underlying distributions that are not true of binomial data or reaction time distributions. It is perhaps wiser to assume that all behavioral time series are *not* stationary (*30*). Therefore we introduce a temporally-local analysis method that compares the reaction times of errors to adjacent correct trials of the same coherence. This analysis cleanly isolates the specific relationship between reaction time and accuracy that is of interest to this study. The temporally-local analysis confirmed our findings in both species: for rats, error trials have shorter reaction times, while for humans errors have longer reaction times (Fig. 4).

Accumulation of evidence (drift diffusion model) is one theoretical framework for understanding reaction times and accuracy in reaction-time perceptual tasks, and our results may have implications for distinguishing among different drift diffusion model variants. But neither our experiment nor our analysis presumes a drift diffusion model, or that evidence is being accumulated, in either species. The data presented here are consistent with evidence accumulation, but we have not done other tests, such as motion kernel analysis (*8*) or choice prediction from reaction time (*31*), to distinguish integration from other strategies, such as extrema detection (*32*).

In this study we considered only visual random dot motion discrimination and only two species. The dependence of accuracy on reaction time likely depends on sensory modality and may even depend on the specific type of visual discrimination. To determine where in the decision-making pathway or when in phylogeny the observed difference arises, more direct species comparisons such as this one are needed, testing other sensory modalities, other types of visual discrimination, and additional species.

Much of the recent theory literature has focused on explaining how accuracy could decline with reaction time, as it does in many human and primate experiments (*1, 25, 26*). Here we present an example in which accuracy increases with reaction time, which is equally demanding of an explanation. Models that account for primate data by positing collapsing decision bounds (*16, 33–37*) or urgency signals (*38–40*) don’t intuitively explain the opposite effect in rat data. The Bayesian optimality argument for collapsing bounds also does not seem consistent with our observation that the decline in accuracy with reaction time in humans is equally prevalent in fixed-coherence epochs. An evidence accumulation model in which the decision time is determined by a soft deadline rather than an evidence threshold could explain an increase in accuracy with reaction time, because decisions which are later are based on more evidence. But there would be no reason to postulate the particular shape of the reaction time distribution we observe, which is characteristic of accumulation to an evidence threshold.

Drift diffusion models that assume noise in the drift rate, starting point, and non-decision time are known to be able to produce either an increase or decrease in accuracy with reaction time (*41*), and thus are at least qualitatively consistent with our data in both species. In particular, variability in the starting point of the diffusion process can produce increasing accuracy with reaction time, whereas variability in drift rate can produce declining accuracy with reaction time. Our subjects do not know a priori which side will be the correct response, so a fixed response preference would manifest as a variable starting point in that model. This could explain why biased rats had stronger positive slopes and biased humans’ slope was less negative, compared to unbiased subjects (Table 1). Fast guessing (ignoring stimulus evidence in a fraction of trials) also produces increasing accuracy with reaction time; this may explain why subjects with >10% lapse (errors on easy trials) had more positive (rat) or less negative (human) slopes. Nevertheless, unbiased rats with <1% lapse show increasing accuracy with reaction time, which is not explained by these phenomena.

It remains unclear to us whether and how other classes of models (*19, 37, 42–45*) might account for both species’ data, but this is an interesting question for future research. It is hoped that exploring this question will lead to model refinements and help distinguish among alternative theories (*46*).

## Methods

### Experimental Methods

#### Ethics

All experiments were performed in strict accordance with all international, federal and local regulations and guidelines for animal and human subject welfare. Experiments involving animals were performed in AAALAC accredited facilities with the approval and under the supervision of the Institutional Animal Care and Use Committee at the University of California, San Diego (IACUC protocol #S04135). This study used 19 Long-Evans rats. Experiments involving human subjects were performed with the approval and under the supervision of the Human Research Protection Program at University of California at San Diego (IRB protocol #141790). This study used 69 healthy adult human volunteers that provided informed consent to enroll in the study.

#### Recruitment and inclusion criteria for human subjects

All human subject data were newly collected for this study. A total of 69 human volunteers were recruited from the UCSD campus by flyers. Both males and females were recruited and no difference between males and females was found. Eligibility criteria for enrollment of the first 14 human subjects (Phase 1 group) were: healthy adult at least 18 years of age, normal or corrected-to-normal vision and ability to hear the feedback tones (all self-reported). Compensation was a flat fee for a 2 hour session, pro-rated for early withdrawal. One subject withdrew after 1 hour. The instructions to subjects evolved over the first 8 subjects, and then remained unchanged (see below). Performance in this cohort was highly variable. Among the subset of subjects that had high trial rates, low lapse (<10%), and fast reaction times, we found that accuracy declined with reaction time; but the results were inconsistent among subjects that were deficient in their effort level (number of trials), lapse, or speed (reaction time).

We therefore recruited a second cohort of 55 new subjects (Phase 2). In an effort to reduce inter-subject variability, the enrollment criteria specified a narrower age range (18-25) and required that the subject was not currently taking any psychiatric medication, in addition to normal or corrected vision and hearing (by self-report). To improve motivation, compensation was linked to performance (number of correct responses, or “points” earned). To provide subjects with performance benchmarks, a leader board was maintained showing the top trial rates and scores. For this cohort, criteria for inclusion of data in analysis were set in advance to require at least 800 trials in 2 hours, with a lapse of less than 10%, and less than 10% of trials with a reaction time >3s (criteria that would have eliminated the “bad” subjects in Phase 1). Eligibility to return for subsequent test sessions was also contingent on satisfying these criteria. All subjects in the second cohort met the performance inclusion criteria for analysis, however, so exclusion was not necessary. Data are shown for all subjects that acquired the task, but we report results separately for Phase 1 and Phase 2 subjects (Table 1). Only Phase 2 subjects were included in the statistical test for positive or negative slope of accuracy vs. reaction time in humans (Methods).

#### Inclusion criteria for rat subjects

Data from 19 Long-Evans rats (Harlan Laboratories) were included in this study. All rats ever tested with random dot motion discrimination task in our lab were included if they successfully acquired the task, were trained and tested with fixed (not ramped) reward size, and had data files containing all the fields required for this analysis. This included 11 male rats that were trained and tested in 2010-2011, and 8 female rats trained and tested in 2016-2017 by entirely different staff in a new location after improvements to the apparatus and overhaul of the code. Five of the male rats also contributed data to Fig. 1c,d and Fig. 2b of (*27*). We note that the rats that were analyzed in detail (Fig. 3-6) in (*27*) were trained and tested with ramped rewards and are not included here. The remaining 6 male and 8 female rats are newly reported in this manuscript. Only the subjects not already reported in (*27*) were included in the statistical test for positive or negative slope of accuracy vs. reaction time (Methods: Statistics).

#### Experimental details common to both species

Experimental trials were delivered on a PC computer (Windows XP-Pro or Windows 7-Pro operating system) using Ratrix (*47*), a custom software suite written in the Matlab environment (MathWorks). Specifically we used Ratrix v1.0.2 and the randomDots stimulus manager. Visual stimuli were rendered on either CRT or LCD monitors using PychToolbox v3.0.8 (*48, 49*). Rendering was synchronized to the 100 Hz vertical refresh of the monitor. Both humans and rats were tested in a darkened room after dark adaptation. Neither rats nor humans were head-fixed or eye-tracked. Auditory feedback was presented through headphones worn on the head (human subjects) or affixed to the left and right sides of training chamber (rats). The times of all subject inputs, visual stimuli, auditory feedback, and water delivery, were recorded in data files.

The trial structure was the same used with rats previously (*27*). A blank screen indicated that the subject could request a trial at any time by a non-rewarded key press (human) or infrared beam break (rat). Upon request without delay, a set of white squares (“dots”) were rendered on the screen at random x,y positions against a black background. At each screen refresh thereafter, each dot was redrawn at a new location that was either random in the x,y plane (noise dots) or displaced by a fixed number of pixels toward the target side (signal dots). Dots that were displaced off the edge of the visual field re-appeared simultaneously on the opposite edge, but with a randomly selected new y position. We verified that the dot density was stably uniform over x and y, the number of coherently moving dots was constant, and spatial patterns among the moving dots continually changed. The motion stimulus ended when the subject responded either L or R. If the response was incorrect, the subject heard a brief error tone and saw a distinct gray screen indicating a time-out, which lasted up to a few seconds before a new trial could be initiated. If the response was correct, the subject immediately heard a brief reward tone; for rats this was paired with water delivery; for humans it was paired with incrementing the number of “points” displayed elsewhere on the monitor. After delivery of reward or timeout, the task returned to the ready screen and waited for a new trial initiation, with no imposed inter-trial interval. This continued until the session was ended by the experimenter.

In most experiments the motion stimulus persisted indefinitely until the subject responded. We found that both humans and rats usually respond within 1-2 seconds; responses later than 3s were rare and were routinely excluded from analysis. In some experiments the moving dots extinguished after 3-5s. If this occurred, the black background persisted until a response was made, with no deadline; responses after the dots extinguished were still rewarded or penalized according to the motion stimulus that had been shown, but responses after stimulus offset were excluded from analysis.

#### Details still differing between human and rat tasks

##### Stimuli

The visual stimuli were chosen to be within a sensitive range for each species, but were not systematically optimized for either species. Rats’ eyes when licking center request port (i.e. at time of stimulus presentation) were at the bottom, centered horizontally, and ~10 cm away from the tangent screen. There were either 100 or 200 dots, displayed at 100% contrast, in a field subtending 125×111 (CRT) or 134×107 (LCD) degrees of visual angle in the upper central visual field. Individual dots were 2-4 degrees on a side, and coherent dots moved at 97 deg/s (CRT) or 73 deg/s (LCD), with 0-100% coherence. The left and right response ports were within the horizontal extent of the moving dot stimulus. Humans’ eyes were centered vertically and horizontally at a distance of ~60cm from an LCD screen. There were more dots (500), displayed at lower contrast (50%) in a smaller field (22 x 13 deg). Individual dots were smaller (0.07 deg), moved slower (28 deg/s), and motion coherence was lower (0-32%).

One human subject was atypical: despite apparent diligence and understanding of the task, their performance was at chance in their first test session. When re-tested on a separate day with a mid-range coherence distribution, performance was above chance and coherence-dependent, with substantial lapse. Upon retesting on a third day with a high-range coherence distribution the subject exhibited high quality psychometric data with<1% lapse, sensitivity 8.8, and threshold coherence 0.37.We have not investigated whether the subjects’ perceptual deficit was specific to motion, or explained by other primary visual deficits in acuity or contrast. This non-representative subject was analyzed separately or excluded in subsequent analyses. This subject’s successful performance was obtained using 100 dots at 100% contrast in a field subtending 43×25 degrees of visual angle. Individual dots were 0.4 degrees on a side, coherent dots moved at 14 deg/s, with 30-100% coherence. We did not parametrically vary parameters other than coherence to determine which of these changes were necessary to rescue performance.

##### Operant modality and reward

Humans were motivated by monetary compensation, indicated trial requests and responses by pressing keys on a keyboard, and were rewarded for correct answers by either “points” (Phase 1) or cash (Phase 2). Rats were motivated by thirst, indicated responses by licking one of two water tubes, and were rewarded for correct answers instantly with water drops. Although humans received monetary rewards only at the end of the two hour session, they received instant auditory reward-predicting feedback at the time of each correct response, and Phase 2 subjects knew that this signal predicted cash reward at the end of the session.

##### Instruction or training

The first few human subjects were required to infer the task rules by trial and error; some failed to acquire the task. After this, verbal instructions with demonstration were developed and remained constant from the 8^th^ subject onward. Subjects were advised that they could take a break at any time. The task began with 50% coherence stimuli and automatically advanced to the testing condition after the subject responded correctly in ten trials in a row. Subjects were eligible to return for up to 5 2-hour testing sessions (in Phase 2 this was subject to minimum performance criteria), but few subjects completed all 5 sessions. Generally we tested two blocks with different coherence distributions in each 2-hour session.

Rats learned the task by operant conditioning through a shaping sequence that lasted for weeks. Briefly, naïve water-restricted rats were initially rewarded for initiating trials at the trial-request port and again for licking the response port below a large salient target stimulus, using large rewards (~200ul), no time limit, and no error penalty. Reward size was gradually reduced (to ~40ul), rewards for trial initiation eliminated, and penalty time-out increased (to 1 s) and this training step continued until rats responded at the visually cued port >80% of trials. Most rats then proceeded directly to the 85% coherence random dot motion task for a learning period of days to weeks until achieving >80% correct performance. (A few rats learned other visual tasks such as image discrimination or orientation discrimination before proceeding to the motion task). Rats acquired proficiency in the motion task over days to weeks.

For rats, during initial training or whenever a response bias developed, correction trials were implemented. After an error trial there was a fixed low probability (≤25%) of entering a “correction trial state”, in which the target response was deterministically the same as the previous trial until the next correct response occurred. The stimulus was still generated randomly each trial, including the selection of coherence and positions of dots. On all other trials, the target side was randomly selected (L or R) with equal probability. Sessions with ≥10% correction trial probability were excluded from analysis. Correction trials were not used with human subjects.

All humans and most rats and were tested in scheduled 2 hour sessions, separated by at least one day. Some rats were also tested in a live-in (24 hour/day access) condition. Most humans acquired task proficiency within the first ~250 trials of their first session, and were tested in only one or two sessions (at most five); we did not study any highly expert psychophysics subjects such as many previous human studies have exclusively used. Rats were tested for dozens to hundreds of sessions. Thus compared to our human subjects, the rats were highly practiced subjects. It will be interesting to learn whether and how highly-practiced human subjects or monkeys differ from our human cohort, but our main goal was to replicate a well-established result in the literature, which we achieved even with comparatively naïve human subjects.

### Data Selection and Analysis Methods

#### Exclusion of learning phase and warm-up trials

Individual testing sessions were included in the analysis only if they contained <10% correction trials and >100 total trials. For purposes of this analysis, the task acquisition criterion for rats was the first occurrence of >80% correct motion direction discrimination over 200 consecutive trials at high coherence; only subsequent training sessions were included in analysis. Subjects that failed to acquire the task to 80% criterion at high coherence were excluded from the study. Human subjects generally performed >80% on high coherence immediately, but the first 250 trials of their first session were excluded as a presumed learning and practice phase.

Both humans and rats tested in 2-hour sessions exhibited “warm-up” effects at the beginning of sessions. Average accuracy improved in the beginning of each session, even in 2^nd^ and 3^rd^ sessions (humans) or after dozens or hundreds of sessions (rats), and even in the absence of any long term trend in accuracy or reaction time over sessions. The average reaction time of rats increased over the beginning part of each session, while humans showed a decrease in reaction time. These trends were often hard to distinguish from chance fluctuations in individual sessions, but apparent after averaging over sessions, and consistent across subjects (not shown).

When rats were tested in 2-hour sessions they were water restricted for the preceding 22 hours. Therefore rushing due to extreme thirst may explain why rats’ reaction times were initially fast and less accurate. When rats were tested in a 24-hour live-in condition, trials were done in bouts which occurred at random times, with weak circadian modulation. The session-onset effect was weaker and perhaps attributable instead to arousal due to handling. This effect stabilized within 100 trials (not shown). We have not analyzed whether within-session bouts exhibited a warm-up effect. Rats typically preformed hundreds of sessions, and thus were highly practiced in the task. Humans, on the other hand, performed only one or a few sessions, so practice effects may explain why reaction time sped up and accuracy increased at the beginning of each session. When the task was stopped and re-started within a 2-hour session, humans showed a weak warm-up effect in the first 10 trials after resuming with new stimulus parameters (not shown).

These warm-up effects were different between species, and are of particular concern because they are in a direction that could cause a spurious positive correlation between reaction time and accuracy for rats, and a negative correlation for humans, potentially explaining the species discrepancy. Therefore we excluded the warm-up trials of every session from our analysis: for rats we excluded the first 200 trials of 2-hour sessions and the first 100 trials of live-in sessions; for humans we excluded the first 250 trials of each session and first 10 trials after resuming after a break within a session.

#### Exclusion of individual trials

Generally there is a clear absolute refractory period of ~500 ms between trial initiation and the earliest responses, which we interpret as reflecting the subject’s non-decision time (visual latency and motor latency). Rarely, shorter reaction times can be caused by false triggering of a lick sensor (such as by a water drop or animal paw), or by a human accidentally hitting two keys at once. To be sure these events were excluded, we took the 1^st^ percentile of each subject’s lifetime reaction times as a conservative estimate of the subject’s non-decision time, and excluded shorter reaction time trials from analysis.

We note that in our previous investigations of speed-accuracy trade-off in rats (*27, 28*) we excluded from analysis all trials that followed error trials. These trials are not typically excluded in the human and primate literature, and were not excluded from any analysis in the present study.

#### Identification of candidate epochs

From each subject’s lifetime data, the trials prior to task acquisition, warm-up trials in each testing session, and trials with reaction times longer than 5 s or shorter than the subject’s lifetime 1^st^ percentile reaction time were excluded from analysis. The subject’s remaining lifetime trials were visualized as concatenated time series. From each contiguous block of trials that had the same reward value, penalty time-out duration, and coherence distribution, we manually selected the trial range that appeared to have stable average accuracy, average reaction time, and qualitative reaction time distribution. In the case of rats, this often entailed combining data from many separate daily sessions. For humans it often entailed selecting a subset of trials from a single session.

The accuracy and reaction times of these candidate epochs were then visualized as smoothed time series, with linear regressions and autocorrelations of both accuracy and reaction time vs. trial number. These plots were compared to shuffled controls in which the temporal order of the trials was randomized. Although fluctuations and trends were often present in the raw time series, these appeared subjectively comparable to the time-shuffled (and thus statistically stationary) controls. All auto-correlograms fell sharply to within shuffle-control levels within ±1 trial and out to ±32 trials (not shown). This screening procedure yielded 144 candidate epochs from 19 unique rats, and 107 candidate epochs from 62 unique human subjects.

We were unable to identify suitable lags for either the Augmented Dickey-Fuller (ADF) test or the Kwiatkowski, Phillips, Schmidt and Shin (KPSS) test for stationarity on the basis of assessing temporally-shuffled controls. This is presumably because neither accuracy nor reaction time were normally distributed variables. Therefore these tests were not used. Instead we used least squares linear regression to estimate the change in the mean of accuracy or reaction time from the beginning to end of the epoch, and a nonparametric correlation (Spearman) to measure the significance of such trends. We considered an epoch to be “trending” if it the correlation between trial number and reaction time involved >50 ms change from beginning to end of the epoch with P<0.01; or >100 ms change, regardless of P value; or if the correlation between trial number and accuracy involved >5% change in accuracy with P<0.01; or >10% change in accuracy regardless of P value. All remaining epochs were designated as “non-trending”. By this criterion there were 120 non-trending epochs from 19 rats, and 87 non-trending epochs from 57 human subjects in our data set.

#### Analysis details for Figure 1

This analysis included any non-trending epoch which included at least 5 coherence values the highest of which was ≥0.8 (for rats or one visually impaired human subject) or ≥0.25 (for humans). A total of N=46 epochs from 11 rats and N=79 epochs from 52 humans qualified for inclusion. The correct (rewarded) response side was assigned randomly each trial; this assignment determined whether or not the response is rewarded or scored as correct, even in a 0% coherence trial. Therefore we expect accuracy to be 50%, and the reaction time distributions of errors and correct trials to be identical, in 0% coherence trials. Psychometric fits were computed using the Palamedes Toolbox (*50*), using the jAPLE and PAL_Logistic options. We constrained the guess rate to 0.5, as this was a 2-alternative forced choice task, and fit the remaining three parameters: lapse, threshold and sensitivity. We defined “high lapse” epochs as those with a lapse of ≥10%, and “low lapse” as all other psychometric epochs. In fixed coherence epochs lapse is not defined. Response bias was evaluated from the number of responses on a side, compared to the number of trials in which that side was the target (correct response). The tendency to respond on side *i* was defined as: 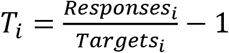, which is 0 if there is no bias, negative if the subject is biased against that side, and positive if biased towards it. Since there were only two response choices, *T*_*L*_ = −*T*_*R*_. We defined “biased” epochs as those with |*T*_*L*_| >=0.1, and other epochs “unbiased”.

#### Analysis details for Figure 2

This analysis included all non-trending, unbiased fixed-coherence epochs, and all non-trending, unbiased, psychometric epochs with ≤10% lapse, excluding the visually impaired human subject. Reaction times longer than 3s were excluded in all panels. A total of 76 epochs from 14 rats and 80 epochs from 55 human subjects qualified for inclusion. For inclusion of a data point in the scatter plots, we further required at least 150 trials of the same coherence occurred within the epoch. Within that set of trials, we determined the average reaction time of the correct vs. error responses, and the accuracy among the fast (bottom quartile of reaction time) vs. slow (top quartile) responses. There were 180 qualifying points for rats and 178 qualifying points for humans.

#### Analysis details for Figure 3

We estimated the slope of the trend of accuracy vs. reaction time for every candidate epoch in the data set that had at least 500 trials. For each epoch, we selected trials with reaction times between the subject’s non-decision time and 3 seconds. If the epoch contained a single coherence, we used all the trials for analysis; if multiple coherences were interleaved, we selected the trials with coherences that had accuracy between 0.7-0.9 based on psychometric fits (c.f. Fig. 1), requiring at least 120 qualifying trials to proceed. We then chose reaction time bin boundaries by quantiles in order to obtain 6 reaction time bins containing equal numbers of trials (≥20 trials/bin). We computed the mean accuracy and mean reaction time in each reaction time bin. We fit a line to reaction time vs. accuracy for all bins that had a mean reaction time <=1.5 seconds, requiring that at least 3 such bins were found (e.g. lines in Fig 3A,D). The slope of this line is the measure reported. Slopes could be estimated in this way for N=111 rat epochs and N=102 human epochs.

For the histograms shown and associated numerical results in text, we included all epochs that were non-trending, unbiased, and had <10% lapse (if psychometric); among these, epochs with <1% lapse, 1-2% lapse, and 2-10% lapse, fixed-coherence epochs, and one visually impaired human subject are distinguished by color in the histograms and broken out in Table 1. We also checked if the results held true in other subsets of the data (Table 1). The predominance of increasing accuracy with reaction time (positive slope) in rats was found regardless of the subset considered: in the data set as a whole (no excluded epochs); in both non-trending than trending epochs; in epochs with either high or low trial numbers; in both psychometric and fixed-coherence experiments; in both biased and unbiased epochs; and in both water-restricted 2-hour sessions and live-in 24-hour sessions.

The predominance of decreasing accuracy with reaction time (negative slope) in humans was found in most subsets, with some exceptions. Negative slopes predominated in the data set as a whole (no exclusions); in both trending and non-trending epochs; both biased and unbiased epochs (though less consistently in the biased subset); and in both Phase 1 and Phase 2 cohorts (though more consistently in Phase 2). One exception was the visually impaired subject, who showed an increase in accuracy with reaction time comparable to rats. This was based on only one non-trending epoch, which was free of lapse or bias. A negative slope was not seen in the subset of human epochs with high lapse rates. This was attributable to positive slopes in two non-trending epochs from the same Phase 2 subject, who also had high bias in both epochs. Slope was negative in human unbiased psychometric epochs of any lapse rate, but near zero on average in the small number of fixed-coherence experiments.

The subset analysis was done to address specific caveats of the analysis, which apply differently to different conditions as mentioned in Discussion. Due to the post-hoc exploratory nature of these regroupings, it is not appropriate to report statistical significance of any of these distinctions. Importantly, however, Table 1 includes every alternative grouping or inclusion criterion that we tested, regardless of the result obtained.

#### Analysis details for Figure 4

The local reaction-time difference between correct and error trials was computed by comparing every error trial to the closest preceding correct trial of the same coherence, enforcing a minimum interval of 3 trials to avoid influence of sequential effects.. These local reaction time differences were then averaged to produce a scalar estimate for any coherence for which at least 50 qualifying pairs of trials were found. We expect this measure to be 0 on average in 0% coherence trials, as correct and error trials are identical from the viewpoint of the subject until after their decision is made.

In Figure 4A, the average local reaction time difference is shown as a function of coherence for a single representative rat epoch. For Figure 4B, local reaction time difference was computed within epoch for N=31 low-lapse psychometric epochs from 7 rats, and then averaged across psychometric epochs; and for N=23 fixed coherence epochs from 9 rats, then averaged over fixed-coherence epochs. For Figure 4C, the local time difference was computed within coherence and then averaged within epoch across all estimated coherences ≥0.40. In the 30 rat psychometric epochs that contained time difference estimates for at least one coherence ≥0.40, the median distance between the compared trials was 34 trials. In the 12 fixed-coherence epochs the median distance was 3 trials.

In Figure 4D, local reaction time difference was computed separately for each coherence based on all the motion coherence task trials performed in one example rat subject’s lifetime with no exclusions. For Figure 4E a lifetime analysis was done separately for N=19 eligible unique rat subjects, and then averaged over subjects. For the histogram in Figure 4F, all 19 subjects had reaction time difference estimates for coherences ≥0.40; the median distance between compared trials was 27 trials.

Analysis of a single human epoch is shown in Figure 4G. For Figure 4H, results from N=66 low-lapse psychometric epochs from 47 subjects were averaged. Results from N=4 fixed coherence epochs from 3 subjects were averaged separately. The histogram in Figure 4I includes the 70 human psychometric epochs for which local reaction time difference estimates were obtained for at least one coherence ≥0.04; the median interval between compared trials in psychometric experiments was 29 trials. All 4 human fixed-coherence epochs used coherence ≥0.04 and are included; among these data the median distance between compared trials was 3 trials.

A single human subject’s lifetime analysis is shown in Figure 4J. Figure 4K is the average of the lifetime analysis from N=69 eligible unique human subjects including one visually impaired human. The histogram in Figure 4L includes all 60 human subjects for which an estimate was obtained for at least one coherence ≥0.04; among these data the median distance between compared trials was 24 trials.

Qualitatively similar results were obtained if instead we compared each error trial to the closest correct trial of the same coherence ≥3 trials later (rather than before).

#### Statistics

We advisedly refrain from assertions of statistical significance in this manuscript (*51*). Recruitment of both rat and human subjects was protracted and open-ended, with no predetermined sample size; data accrual was terminated for reasons of personnel departure and funding exhaustion. The analysis was ongoing during data collection and afterwards, and analysis methods were highly exploratory and iteratively refined. In particular, we tried many different ways of detecting and excluding nonstationary data; many different criteria for including or excluding subjects, sessions or trials; and numerous ways of analyzing for the relationship between reaction time and accuracy. Project progress reports included interim statistical analysis. Because it would be impossible to enumerate the number of implicit tests performed, multiple comparison correction is not feasible, and therefore P values are generally not stated. With the exception of the preliminary results from Phase I human subjects (discussed above), however, we note that we obtained the same qualitative results at every phase of interim analysis and with every alternative analysis method we tried. Accrual of additional subjects and refinement of analysis methods were aimed at increasing stringency to determine if the positive result already obtained could be falsified. After procedures for selecting candidate epochs and excluding trials were finalized as described above, every alternative subset analysis (regrouping) that we tested is reported regardless of outcome (Table 1).

After completing all other analysis in this manuscript, we computed P values for the main claim of the paper, once only, as follows. We reasoned that the most valid data to use were the non-trending, unbiased, low-lapse (<10% lapse) epochs. We included only the human subjects recruited in Phase 2, for which inclusion criteria had been established prior to recruitment (but among which none were excluded by those criteria). The visually impaired subject was not in this group. To ensure independence of statistics in separate reports, only the rat subjects that were not already reported in (*27*) were included here. We used the measure of slope of accuracy vs. reaction time (c.f. Fig. 3 C,F) as the measure of the effect. Within each subject, if a slope had been determined for more than one epoch, we used the average of the slopes, such that samples were independent (only one value per subject). We then performed the nonparametric Wilcoxon Sign Rank test against the null hypothesis that the median of the population is >0 or <0 respectively. From this analysis we obtained: Human population slope −0.18±0.18 mean±SD, N=43 subjects, P(>0)=1.00, P(<0)=1.58e-07. Rat population slope 0.18±0.09, N=9 subjects, P(>0)=1.95e-03, P(<0)=1.00.

## Author contributions

CAS co-designed and implemented human experiments, recruited and tested human subjects, and performed the interim analysis similar to Figures 1–3 for a subset of the human subjects. PR conceived of the study, designed and implemented the rat experiments, co-designed human experiments, provided oversight of all data collection, performed all analysis in the manuscript, made the figures, and wrote the manuscript. The authors declare no conflicts of interest.

## Acknowledgements

In a technical capacity, CAS provided expert support of all rat experiments. Serena Park, Ryan Makin, Nicole Dones, Anjali Herekar, Michaela Juels, Xiao Guo, Vishal Venkatraman and Grace Lo helped provide daily health monitoring of rats and ran behavioral testing sessions. Michaela Juels helped recruit and test human subjects. Support for this research was provided by a Kavli Institute for Brain and Mind Innovative Research Grant; the Regents of California Academic Senate General Campus Research and Bridge Fund programs; and a UCSD Division of Biology Bridge Grant. This manuscript has been released as a preprint at bioRxiv (*52*).

